# New insights on the pharmacogenomics of antidepressant response from the GENDEP and STAR*D studies: rare variant analysis and high-density imputation

**DOI:** 10.1101/109827

**Authors:** Chiara Fabbri, Katherine E. Tansey, Roy H. Perlis, Joanna Hauser, Neven Henigsberg, Wolfgang Maier, Ole Mors, Anna Placentino, Marcella Rietschel, Daniel Souery, Gerome Breen, Charles Curtis, Lee Sang-Hyuk, Stephen Newhouse, Hamel Patel, Michel Guipponi, Nader Perroud, Guido Bondolfi, Micheal O’Donovan, Glyn Lewis, Joanna M. Biernacka, Richard M. Weinshilboum, Anne Farmer, Katherine J. Aitchison, Ian Craig, Peter McGuffin, Rudolf Uher, Cathryn M. Lewis

## Abstract

Genome-wide association studies have generally failed to identify polymorphisms associated with antidepressant response. Possible reasons include limited coverage of genetic variants that this study tried to address by exome genotyping and dense imputation.

A meta-analysis of Genome-Based Therapeutic Drugs for Depression (GENDEP) and Sequenced Treatment Alternatives to Relieve Depression (STAR*D) studies was performed at SNP, gene and pathway level. Coverage of genetic variants was increased compared to previous studies by adding exome genotypes to previously available genome-wide data and using the Haplotype Reference Consortium panel for imputation. Standard quality control was applied. Phenotypes were symptom improvement and remission after 12 weeks of antidepressant treatment. NEWMEDS consortium samples and Pharmacogenomic Research Network Antidepressant Medication Pharmacogenomic Study (PGRN-AMPS) served for replication.

7,062,950 SNPs were analysed in GENDEP (n=738) and STAR*D (n=1409). rs116692768 (p=1.80e-08, *ITGA9* (integrin alpha 9)) and rs76191705 (p=2.59e-08, *NRXN3* (neurexin 3)) were significantly associated with symptom improvement during citalopram/escitalopram treatment. At gene level, no consistent effect was found. At pathway level, the Gene Ontology terms GO:0005694 (chromosome) and GO:0044427 (chromosomal part) were associated with improvement (corrected p=0.007 and 0.045, respectively). The association between rs116692768 and symptom improvement was replicated in PGRN-AMPS (p=0.047), while rs76191705 was not. The two SNPs did not replicate in NEWMEDS.

*ITGA9* codes for a membrane receptor for neurotrophins and *NRXN3* is a transmembrane neuronal adhesion receptor involved in synaptic differentiation. Despite their meaningful biological rationale for being involved in antidepressant effect, no convincing replication was achieved. Further studies may help in clarifying their role.

## 1. Introduction

Major depressive disorder (MDD) became one of the five leading diseases contributing to disability-adjusted life years (DALYs) in 2010 in the US (1). MDD is associated with a huge increase in suicide risk (2), poor quality of life (comparable to that observed in severe physical disorders such as arthritis and heart disease (3)) and health expenditure (direct costs alone amount to 42 billion dollars per year in Europe (4)).

Despite the availability of antidepressant drugs belonging to different classes, high inter-individual variability is observed in response. The lack of reliable and reproducible markers of treatment outcome contributes to unsatisfactory response and remission rates as well as to side effect burden, poor treatment adherence and early treatment discontinuation (5). Following the observation that antidepressant response clusters in families, genetic variants were considered promising biomarkers to tailor antidepressant treatments and improve the prognosis of MDD (6, 7). Genome-wide association studies (GWAS) were a promising tool to identify the polymorphisms involved in antidepressant response after the overall contradictory and non-replicated findings of candidate gene studies (8). But GWAS results fell below expectations, with no genome-wide significant signal (p<5e-08) that was replicated in different samples (9–15). Possible reasons for these disappointing results include: 1) limited coverage of genetic variants (e.g. ~ 500 K common polymorphisms were originally analyzed in STAR*D, GENDEP and MARS studies and ~ 1.2 millions in the meta-analysis of these studies thanks to imputation, while ~ 40 millions polymorphisms are known to date thanks to sequencing studies (9–11) (12, 16); 2) limited sample size; 3) sample heterogeneity (e.g. different subtypes of depression and severity, different antidepressants); 4) analysis of common variants (minor allele frequency (MAF) > 0.01) alone. Further, previous GWAS meta-analyses focused on single marker analysis and pathway analysis was performed in single samples and not as meta-analysis among different samples (12, 17).

Considering these limitations, the current study aimed to:

1. Increase the coverage of genetic variants by analysing exonic polymorphisms and dense imputation;
2. Test for association with antidepressant response at the level of SNPs, genes and pathways including both common and rare variants;
3. Reduce heterogeneity in treatment by analysing patients treated with the same antidepressant, as performed previously (e.g. (10, 17)).

In addition to the discovery samples of GENDEP and STAR*D, two replication samples were available (NEWMEDS and PGRN-AMPS).

## 2. Materials and methods

### 2.1 Samples

#### 2.1.1 GENDEP

The Genome-Based Therapeutic Drugs for Depression (GENDEP) project was a 12-week partially randomized open-label pharmacogenetic study with two active treatment arms. 867 patients with unipolar depression (ICD-10 or DSM-IV criteria) aged 19–72 years were recruited at nine European centres. Eligible participants were allocated to flexible-dosage treatment with either escitalopram (10–30 mg daily, 499 subjects) or nortriptyline (50–150 mg daily, 368 subjects). Severity of depression was assessed weekly by the Montgomery-Asberg Depression Rating Scale (MADRS) (18), Hamilton Rating Scale for Depression (HRSD–17) (19) and Beck Depression Inventory (BDI) (20). Detailed information about the GENDEP study has been previously reported (10).

#### 2.1.2 STAR*D

The Sequenced Treatment Alternatives to Relieve Depression (STAR*D) study was a NIMH-funded study aimed to determine the effectiveness of different treatments for patients with MDD who have not responded to the first antidepressant treatment. Non-psychotic MDD (DSM-IV criteria) patients with age between 18 and 75 years were enrolled from primary care or psychiatric outpatient clinics. Severity of depression was assessed using the 16-item Quick Inventory of Depressive Symptomatology-Clinician Rated (QIDS-C16) (21) at baseline, weeks 2, 4, 6, 9, and 12, while HRSD–17 was administered at each level entry and exit. All patients received citalopram in level 1 and the present study is based on level 1 data. 1953 patients were included in the genetic study. Detailed description of the study design and population are reported elsewhere (22).

#### 2.1.3 Replication samples

NEWMEDS consortium (http://www.newmeds-europe.com) (23) samples other than GENDEP (17) and PGRN-AMPS (Pharmacogenomic Research Network Antidepressant Medication Pharmacogenomic Study) sample (14) were used for replication of significant findings obtained in the GENDEP-STAR*D meta-analysis.

As part of the NEWMEDS consortium (for further details see (17)), three studies conducted by academic institutions (GENDEP, see paragraph 2.1.1; GENPOD, a randomized controlled trial of two active antidepressants, n □=□ 601 (24); and GODS, a treatment cohort of severe depression, n □=□ 131 (25)) and two studies by pharmaceutical industry members of the European Federation of Pharmaceutical Industries and Associations (active comparator arms from randomized controlled trials by Pfizer, n □=□ 355, and GlaxoSmithKline, n □=□ 191) were combined. All included patients were diagnosed with MDD and treated for 6 to 12 weeks with either an antidepressant that acts primarily through blocking the reuptake of serotonin (SSRIs: escitalopram, citalopram, paroxetine, sertraline, fluoxetine) or an antidepressant that acts primarily through blocking the reuptake of norepinephrine (NRIs: nortriptyline, reboxetine), see (17) for details.

PGRN-AMPS included 529 participants with nonpsychotic MDD who were recruited at the Mayo Clinic in Rochester, Minnesota primarily through the inpatient and outpatient practices of the Department of Psychiatry and Psychology. Participants were offered an eight-week course of treatment with either citalopram or escitalopram and depressive symptoms were rated using QIDS-C16 as in the STAR*D study in addition to the HRSD–17. For further details see (14).

### 2.2 Outcomes

The primary outcome of this study was depressive symptom improvement after 12 weeks of antidepressant treatment. Continuous measures, such as percentage improvement, capture more information and have higher power than cutoff-based dichotomous measures, such as remission, however remission was associated with MDD prognosis (26) (27). The percentage change in scores between baseline and 12 weeks was used to measure symptom improvement, using MADRS in the GENDEP study, and QIDS-C16 in STAR*D, as in previous studies in these samples (10, 28).

As a secondary outcome, we investigated symptom remission after 12 weeks of antidepressant treatment. According to standard criteria, remission was defined as HRSD–17 ≤ 7 (29) and QIDS-C16 ≤ 5 (28) in GENDEP and STAR*D, respectively. HRSD–17 was used to define remission in the GENDEP given the stronger consensus about the threshold to identify remission on this scale in contrast to MADRS, where different definitions of remission have been reported (≤ 12 (30), ≤ 10 (31), ≤ 8 (32)).

Each outcome measure (percentage change, remission) was analyzed separately in GENDEP and STAR*D, and then a meta-analysis performed. Two analyses were performed, initially using all samples, and then including only escitalopram-treated patients from GENDEP, since escitalopram is the active isomer of citalopram (33), the antidepressant used in STAR*D level 1.

Missing data were handled as in previous studies on the investigated samples (9, 10). When at least one post-baseline assessment was available, the percentage improvement at 12 weeks was estimated as the best unbiased estimate of mixed effect linear models. Participants without any post-baseline measurement were excluded from the analyses. Since specific antidepressant response is associated with depression severity (34), a minimum depression severity score of 14 on the HRSD–17 was an inclusion criterion in the STAR*D study (all the subjects that we included satisfied this criteria) but not in GENDEP thus a sensitivity analysis excluding GENDEP subjects with HRSD–17<14 was performed for the validation of significant findings.

### 2.3. Genotyping and imputation

Genome-wide data available in STAR*D were obtained using Affymetrix Human Mapping 500K Array Set in 969 subjects and Affymetrix Genome-Wide Human SNP Array 5.0 (Affymetrix, South San Francisco, California) in the remaining 979 samples, while in GENDEP Illumina Human610-quad bead chip (Illumina, Inc., San Diego) was used (9, 10). Further genotyping in both samples was performed by the Illumina Infinium Exome-24 v1.0 BeadChip that includes ~ 250K variants. Pre-imputation quality control was performed according to the following criteria: 1) variants with missing rate ≥ 5%; 2) monomorphic variants; 3) subjects with genotyping rate < 97%; 4) subjects with gender discrepancies; 5) subjects with abnormal heterozygosity; 6) related subjects (identity by descent (IBD) >0.1875 (35)); 7) population outliers according to Eigensoft analysis of linkage-disequilibrium-pruned genetic data (36, 37); 8) GWAS discordant subjects (referred to exome data only) and 9) non-white subjects (referred to STAR*D only since all subjects included in the GENDEP are of Caucasian ethnicity). Hardy–Weinberg equilibrium was not used as an exclusion criterion for markers (as previously done in the same datasets (17)), since departures from Hardy– Weinberg equilibrium are expected in a case-only study (38).

Data were imputed using Minimac3 as provided by the Michigan imputation Server (https://imputationserver.sph.umich.edu/start.html). Post-imputation quality control was performed pruning variants according to the following criteria: 1) poor imputation quality (R^2^ < 0.30 (39, 40); 2) minor allele frequency (MAF) < 0.01 (see further details in paragraph 2.4 Statistical analysis). For a flow chart describing pre-and post-imputation quality control on each dataset see Supplementary Figure 1. Since exome array data were available only in 1015 subjects from the STAR*D sample (after quality control), the imputation of these data was performed separately from the imputation of genome-wide array data (1470 subjects).

**Figure 1:**
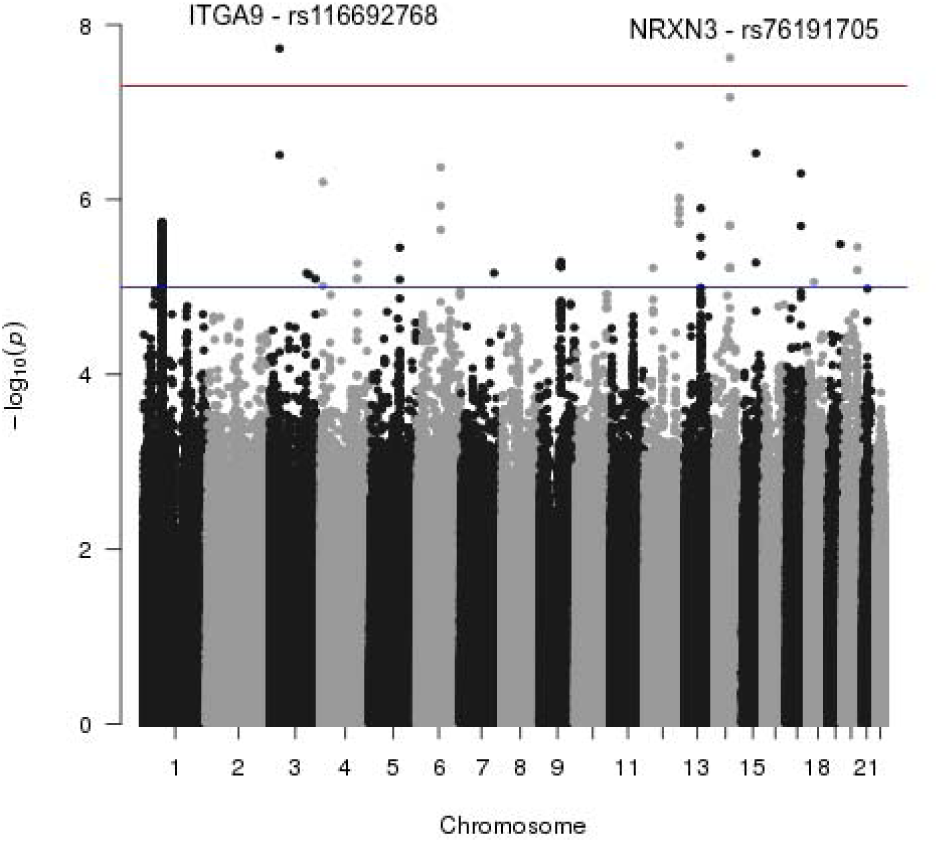
Manhattan plot referred to symptom improvement in STAR*D and GENDEP escitalopram-treated subsample meta-analysis.

### 2.4 Statistical analysis

We performed a fixed-effects meta-analysis to test the association between single polymorphisms and phenotypes using PLINK (41). Heterogeneity measures (Cochrane’s Q statistic and the I^2^ heterogeneity index) were calculated. We tested linear or logistic regression models including the ancestry-informative principal components, recruitment centre, age, and baseline severity as covariates in line with previous publications on these samples (10, 12, 42). The same covariates were used for gene and pathway analysis.

We tested the association between genes and phenotypes as well as pathways and phenotypes using MAGMA (43). MAGMA performs both a self-contained and a competitive gene-set analysis, the latter is more conservative and it was applied in this study since it reflects the difference in association between genes in the pathway and genes outside the pathway. Only for the replication analyses self-contained analysis results were also reported since the replication nature of these tests.

Both rare (MAF<0.01) and common variants were included in gene and pathway analysis, but only genotyped rare variants were retained while imputed rare variants were excluded. Indeed imputation quality of rare variants using the HRC panel was found to be better than using 1000 Genomes data but still not great (44). Thus we chose a relatively conservative approach, including only subjects genotyped on both genome-wide and exome array in the gene and pathway analysis. In the gene- and pathway-level meta-analysis different weights were assigned to polymorphisms according to their MAF, thus higher weight was assigned to rare variants as implemented in MAGMA (43).

For pathway analysis in each dataset and their meta-analysis the reported results refer to a competitive gene-set analysis, which uses a conditional model to correct for confounding due to gene size, gene density and (if applicable) differences in sample size per gene (43). The analysed pathways were downloaded from http://software.broadinstitute.org/gsea/downloads.jsp (Biocarta, KEGG, Gene Ontology, Reactome, microRNA targets and transcription factor targets).

An attempt to replicate significant results was performed in NEWMEDS (omitting GENDEP) and PGRN-AMPS. In NEWMEDS phenotypes and covariates were as described in a previous study (17). Briefly, percent symptom improvement at end-point was adjusted for covariates (age, gender, baseline severity, ancestry-informative principal components and centre in case of multi-centric studies) and z-score transformed. Samples were genotyped on Illumina Human610-Quad BeadChips or Illumina Human660W-Quad BeadChips.

For PGRN-AMPS details about phenotypes and covariates were described elsewhere (14). Percent symptom improvement at end-point was used as phenotype and covariates were the first four population principal components and age as in the original GWAS (14). Samples were genotyped on Illumina Human610-Quad BeadChips (Illumina, San Diego, CA).

In both replication samples we performed genotype imputation using the same method applied in GENDEP and STAR*D; pre- and post-imputation quality control were performed according to the same criteria. For replication of individual SNP results, the index SNP and those in linkage disequilibrium (R^2^≥0.30) were considered.

### 2.5 Multiple-testing correction and power analysis

For individual SNP analysis, a genome-wide significance threshold was set at p=5e-08. A suggestive significance threshold was set at a p value of 5e-06, which is two orders of magnitude below the genome-wide significance level and approximately corresponds to a level at which one association per genome-wide analysis is expected by chance (45).

For MAGMA gene level analysis, the False Discovery Rate (FDR) correction was applied. For MAGMA pathway analysis, 50,000 permutations were performed to correct for multiple testing.

For a continuous outcome in the whole sample (n=2145) and setting alpha=5e-08, we had 80% power to identify a SNP with an effect size (heritability) of 0.018, while in citalopram-escitalopram treated sample (n=1739) we had 80% power to identify an effect size of 0.022 (46).

For a dichotomous phenotype in the whole sample and setting alpha=5e-08, we had 80% power to identify a risk allele with MAF=0.06 and RR=1.50, while in citalopram-escitalopram treated sample we had 80% power to identify a risk allele with MAF=0.07 and RR=1.50 (47).

## 3. Results

The clinical-demographic characteristics of the samples are reported in Supplementary Table 1. 2145 subjects (1409 and 736 from STAR*D and GENDEP, respectively) were included in the SNP-level meta-analysis, 1828 of them were treated with citalopram or escitalopram. 1739 (1003 and 736 from STAR*D and GENDEP, respectively) subjects were included in the gene and pathway meta-analysis, 1422 of them were treated with citalopram or escitalopram.

**Table 1:** SNPs showing genome-wide association with symptom improvement in GENDEP escitalopram-treated sample and STAR*D meta-analysis. Chr=chromosome. Pos=position (GRCh37). NMD=non-sense mediated decay. Q=Cochrane’s Q statistic. I=I^2^ heterogeneity index. Rsq=measure of imputation quality. MAF=minor allele frequency. Random and fixed p values corresponded for these SNPs.

| **SNP** | **Chr** | **Base pair** | **Gene** | **P** | **Q** | **I** | **Rsq STARD/escGENDEP** | **Beta STARD/escGENDEP** | **MAF STARD /escGENDEP** |
| --- | --- | --- | --- | --- | --- | --- | --- | --- | --- |
| s116692768 | 3 | 37540482 | *ITGA9*(intron) | 1.87e-08 | 0.36 | 0 | 0.75 / 0.78 | -0.14 / -0.20 | 0.035 / 0.023 |
| s76191705 | 14 | 79718462 | *NRXN3*(intron) | 2.39e-08 | 0.76 | 0 | 0.61 / 0.96 | -0.26 / -0.23 | 0.010 / 0.016 |

### 3.1 SNP analysis results

The GENDEP and STAR*D meta-analysis included 7,062,950 SNPs and showed no evidence of genomic inflation (lambda values were ≤ 1.01, QQ-plots are shown in Supplementary Figure 2).

**Figure 2:**
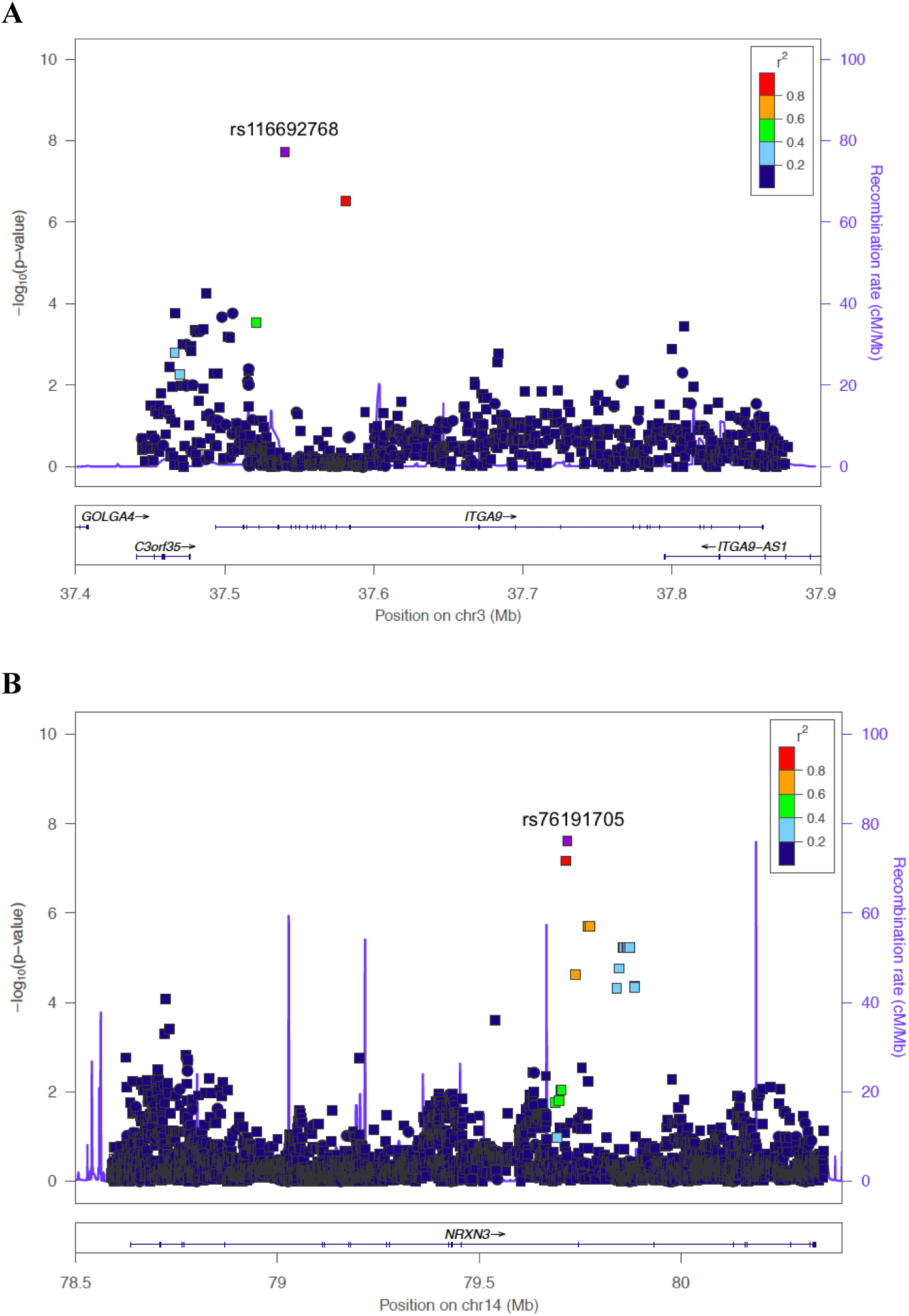
regional association plots referred to ITGA9 (**A**) and NRXN3 (**B**) and symptom improvement in STAR*D and GENDEP escitalopram-treated subsample. Plots were obtained using LocusZoom (PMID 20634204). Imputed SNP are plotted as squares and genotyped (in at least one sample) SNPs as circles.

In the whole sample, no SNP reached the genome-wide significance threshold for association with symptom improvement or remission. Eighty SNPs, from 17 genomic regions, reached suggestive level of association (p<5e-06) (Supplementary Table 2).

In STAR*D and escGENDEP (GENDEP escitalopram) meta-analysis, rs116692768 (MAF=0.033, beta STAR*D=0.14, beta escGENDEP=-0.20, p=1.87e-08) and rs76191705 (MAF=0.012, beta STAR*D=-0.26, beta escGENDEP=-0.23, p=2.39e-08) were significantly associated with symptom improvement (Figure 1; Table 1). These SNPs are located within introns of the *ITGA9* (integrin alpha 9) and *NRXN3* (neurexin 3) genes. Another *NRXN3* SNP (rs79302561) was close to the significance threshold (p=6.76e-08). There was no heterogeneity as measured by I^2^ for these three SNPs, thus fixed-effects and random-effects p values corresponded. Each SNP was imputed, with R^2^ values of over 0.6 for each SNP in each cohort (Table 1). These SNPs retained significance after the exclusion of GENDEP patients with baseline HRSD–17<14 (p=2.69e-08 and 3.43e-08, respectively). Regional association plots for *ITGA9* and *NRXN3* genes are reported in Figure 2. An overview of SNPs with suggestive level of association (p<5e-06) with symptom improvement in STAR*D and escGENDEP is reported in Supplementary Table 3.

In the analysis of remission, no SNPs reached significance in the full meta-analysis or the meta-analysis with escGENDEP, SNPs with suggestive level of association (p<5e-06) are reported in Supplementary Tables 4-5.

### 3.2 Gene analysis results

The gene-level analysis included 17,996 genes with 3,185,134 SNPs.

In the whole sample meta-analysis, the olfactory receptor family 4 subfamily K member 2 (*OR4K2*) gene was associated with symptom improvement after multiple-testing correction (nominal p=2.13e-06, corrected p=0.038 (FDR)). *OR4K2* included 4 rare genotyped SNPs in both datasets but the overall effect of *OR4K2* rare alleles on improvement was in the opposite direction between the samples. GENDEP subjects carrying rare alleles (rs199718838 A, rs116972349 A, rs151057533 C and rs147651981 T, n=8 subjects) showed lower mean symptom improvement (18.58±53.43%) compared to common alleles carriers (59.78±24.49%), while in STAR*D the opposite was found (fourteen subjects carrying rare alleles (rs199718838 A, rs116972349 A, rs142549715 A and rs150417989 G) had a mean improvement of 70.30±22.23% compared to a mean improvement of 50.33±32.98% in common allele carriers). No gene survived multiple-testing correction in the analysis of remission (Supplementary Table 6).

In the meta-analysis of STAR*D and escGENDEP, no gene was associated with symptom improvement or remission (Supplementary Table 7). Several genes with nominal p≤0.0005 overlapped with those found in the whole sample, such as *POU1F1, PAG1, PKM, RPUSD3* and *PARP6.*

### 3.3 Pathway analysis results

17,996 genes including 3,185,134 SNPs were included in this analysis.

In the whole sample, the Gene Ontology term corresponding to the chromosome pathway (GO:0005694) and the chromosomal part (GO:0044427) pathway were associated with symptom improvement (corrected p=0.007 and 0.045, respectively). No pathway was associated with remission. The pathways including genes with promoter regions around transcription start site containing the motif TGACGTYA which matches annotation for JUN oncogene and genes with promoter regions around transcription start site containing the motif TGACGTMA which matches annotation for CREB1 showed a trend of association (corrected p=0.092 and 0.094, respectively).

In STAR*D and escGENDEP, the strongest evidence of association was obtained from the steroid hormone receptor signaling pathway (GO:0030518) and the intracellular receptor mediated signaling pathway (GO:0030522) for symptom improvement (corrected p=0.070 and 0.087, respectively) and remission (corrected p=0.055 and 0.079, respectively).

An overview of results is reported in Table 2 and the functional role of the variants in each of these pathways is shown in Supplementary Figure 3. Interestingly, the chromosome pathway (GO:0005694) was the richest in rare missense variants (6.6% versus 2.5-3% in the other pathways).

**Table 2:** results of pathway meta-analysis in the whole sample and STAR*D – GENDEP escitalopram subgroup (CIT-ESCI). Results with corrected (permutated) p < 0.10 are reported. For each pathway “Top genes” are those showing nominal p ≤ 0.10 for association with the phenotype.

| Pathway | Sample Phenotype | N genes | Top genes | P |
| --- | --- | --- | --- | --- |
| Chromosome pathway (GO:0005694) | Whole sample - improvement | 116 | NPM2, PAM, BUB1, UPF1, POLG2, NDC80, RFC4, HMGB1, FOXC1, TIPIN, SMC2, TOP1, H2AFY, OIP5 | 0.0069 |
| Chromosomal part (GO:0044427) |  | 92 | NPM2, PAM, BUB1, UPF1, NDC80, RFC4, FOXC1, SMC2, TIPIN, OIP5, H2AFY | 0.045 |
| Targets of JUN |  | 247 | CCNA2, GPBP1, VGF, GNB4, RAI1, RUNDC3A, AHI1, UBE2H, ALS2, ABHD16A, SIK1, SLC18A2, PDP1, PCSK1, IRF2BPL, ELAVL1, ELOVL5, ANAPC10, TIPRL, B3GALT2, HOXC10, RPS29, ZFAND2B, SYNCRIP, MAP3K13, LGR5, RAB25, TSC22D2, PACRGL, LENG9, GLI1, TGIF2, GPR3, RCE1, ATG5, TRPC1, HHIP, SYT11, RBM18 | 0.092 |
| Targets of CREB1 |  | 250 | CCNA2, GPBP1, VGF, GNB4, RAI1, RUNDC3A, AHI1, UBE2H, ALS2, ABHD16A, SIK1, SLC18A2, PDP1, PCSK1, IRF2BPL, ELAVL1, ELOVL5, ANAPC10, TIPRL, HOXC10, RPS29, ZFAND2B, MAP3K13, RBMS2, LGR5, RAB25, TSC22D2, ZNF687, TUBB2B, PACRGL, GLI1, TGIF2, GPR3, RCE1, ATG5, HHIP, SYT11, RBM18 | 0.094 |
| Steroid hormone receptor signaling pathway (GO:0030518) | CIT-ESCI – improvement | 18 | UBR5 | 0.070 |
| Intracellular receptor mediated signaling pathway (GO:0030522) |  | 19 | UBR5 | 0.087 |
| Steroid hormone receptor signaling pathway (GO:0030518) | CIT-ESCI remission | 18 | YWHAH, MED13, DAXX | 0.055 |
| Intracellular receptor mediated signaling pathway (GO:0030522) |  | 19 | YWHAH, MED13, DAXX | 0.079 |

### 3.4 Replication samples

In both replication samples, only the genome-wide significant findings of STAR*D–escGENDEP meta-analysis and independent SNPs showing R^2^≥0.30 with them were analysed. Secondly, pathways associated with phenotypes in STAR*D-GENDEP meta-analysis were also investigated. 894 and 492 subjects were included after quality control in NEWMEDS and PGRN-AMPS, respectively.

In NEWMEDS the two SNPs associated with symptom improvement in the STAR*D – escGENDEP meta-analysis were not replicated and weak nominal associations were found for two SNPs (s7152916 and rs7152941) in LD with *NRXN3* rs76191705 (Supplementary Table 8A). In this sample only three independent SNPs in LD with the index SNPs were available.

Over 90% of SNPs included in the pathways showing significant associations (or trends, see Table 2) in the GENDEP-STAR*D meta-analysis were available also in NEWMEDS and only these pathways were analysed in NEWMEDS. The chromosome and chromosomal part pathways were not associated with symptom improvement (nominal p=0.34 and 0.45, respectively) nor were pathways including targets of JUN and targets of CREB1 (nominal p=0.78 and 0.95, respectively). In the subsample treated with citalopram or escitalopram (n=370), the intracellular receptor mediated signaling pathway and the steroid hormone receptor signaling pathway were associated with improvement (permutated comparative p=0.0028 and p=0.017, respectively, self-p=0.0089 and 0.014 (self-p values are also reported because only six pathways were analysed here and for a replication purpose, see paragraph 2.4 Statistical analysis for details about comparative and self-p values)) but not remission (p=0.11 and 0.08, respectively).

In PGRN-AMPS only *ITGA9* rs116692768 and *NRXN3* rs76191705 were investigated because no SNP showed LD≥0.30 with them. rs116692768 was associated with symptom improvement in the same direction to STAR*D – escGENDEP meta-analysis (p=0.047, Supplementary Table 8B; no multiple testing correction was applied here since only two SNPs were investigated for replication purpose), while rs76191705 showed no association with this phenotype (p=0.34).

In PGRN-AMPS only the intracellular receptor mediated signaling pathway and the steroid hormone receptor signaling pathway were analysed because all patients were treated with citalopram or escitalopram in this sample. Over 90% of SNPs included in these pathways in STAR*D – escGENDEP meta-analysis was available in PGRN-AMPS, but no association with symptom improvement was found (nominal self-p values were 0.10 and 0.11, respectively).

## 4. Discussion

### 4.1 Main findings

The present study has exploited existing pharmacogenetic samples (GENDEP and STAR*D), through: 1) the increase of genetic variant coverage (exome genotyping, high-density imputation) and 2) SNP-, gene- and pathway-level meta-analysis.

In the SNP-level meta-analysis, rs116692768 and rs76191705 were associated with symptom improvement during citalopram/escitalopram treatment (Table 1), rs116692768 was nominally replicated in PGRN-AMPS but none of the two SNPs was replicated in the remaining NEWMEDS samples. These SNPs are located within *ITGA9* (integrin subunit alpha 9) *NRXN3* (neurexin 3), respectively. Both these SNPs are intronic, but the latter is a NMD (non-sense mediated decay) transcript variant (i.e. it is a target of NMD). NMD is a post-transcriptional surveillance process that recognizes and degrades mRNAs containing premature termination codons but it also targets 3– 10% of normal transcripts in humans, thereby serving as a widespread gene regulatory mechanism (48–50). rs79302561 T allele was reported to act as an enhancer of gene expression in several cell types (Ensembl GRCh37 release 84). *ITGA9* rs143661452 lies within both *ITGA9* and *ITGA9*-AS1 (antisense RNA). Antisense RNAs are involved in multiple regulatory processes in eukaryotes, such as transcriptional interference and RNA masking (51). No regulatory role of rs143661452 is known so far.

The proteins coded by *ITGA9* and *NRXN3* show interesting functions in relation to antidepressant action. Integrins are heterodimeric (one alpha and one beta subunit) transmembrane proteins that connect the extracellular environment to intracellular signaling. In the CNS they are involved in the control of synaptic plasticity, long-term potentiation (LTP), cell adhesion and migration (52, 53). Polymorphisms in another beta isoform (*ITGB3*) and expression level of this gene have been associated with antidepressant response in humans (42, 54). Neurexins are type I transmembrane neuronal adhesion receptors and their interaction with neuroligins is sufficient to trigger postsynaptic and presynaptic differentiation (55).

Non-genome-wide significant SNPs showing suggestive p-values (<5e-06) were mainly intergenic but several of them lie in genes previously reported to be involved in mood disorders or antidepressant response (particularly *SORCS2, GRIN2D, CTNND2, CSGALNACT1, DISC1, TSNAX-DISC1*) (56–65).

The gene-level meta-analysis did not show any consistent finding. Indeed the olfactory receptor family 4 subfamily K member 2 (*OR4K2*) gene reached the significance threshold (Supplementary Table 6) but the effect of rare alleles on symptom improvement had opposite direction between GENDEP and STAR*D. Olfactory receptor genes have been shown to be over-represented among homozygous loss of function (HLOF) genes and segregating polymorphisms of functional and non-functional copies of olfactory genes are common (66). The olfactory receptor family is the most polymorphic family of genes in humans after the major histocompatibility complex and the phenotypic consequences of such genetic variability are not completely clear but likely not dramatic given their large diffusion (67).

Our pathway meta-analysis in GENDEP-STAR*D showed that the chromosome pathway (GO:0005694) and the chromosomal part (GO:0044427) pathway were associated with symptom improvement. These pathways were not associated with improvement in NEWMEDS, but GO:0044427 and particularly GO:0005694 were much richer in rare missense variants compared to the other significant pathways (Supplementary Figure 3) and this may have limited the replicability of the finding since no exome genotyping was performed in NEWMEDS. Interestingly, the chromosomal part pathway has been previously associated with antidepressant response by a genome-wide gene expression study (68). Among the top genes of the chromosome and chromosomal part pathways (Table 2), some are involved in the differentiation of neural stem cells into neurons (*UPF1* (Regulator Of Nonsense Transcripts Homolog), HMGB1 (High Mobility Group Box 1) and neural development (*FOXC1* (Forkhead Box C1) (69–71)). Thus these genes may play a role in the neurogenesis process that is known to mediate the effect of antidepressants. Another member of these pathways is *PAM* (Peptidylglycine Alpha-Amidating Monooxygenase) that acts as a regulator of amygdala excitability and synaptic plasticity (72) and consistently it plays a role in emotional responses regulation (73).

The meta-analysis of escGENDEP and STAR*D showed that the steroid hormone receptor signaling pathway and the intracellular receptor mediated signaling pathway (that largely overlap between each other) were close to the significance threshold after permutations. Impaired glucocorticoid receptor (GR) function has been suggested to be causal for HPA axis hyperactivity in MDD that results in impaired neurogenesis and reduction of hippocampal volume. Antidepressants modulate the expression of GR, its translocation to the nucleus and also the transcription of its target genes (74). The steroid hormone receptor signaling pathway and the intracellular receptor mediated signaling pathway were associated with improvement in NEWMEDS citalopram-escitalopram treated subsample but not in PGRN-AMPS. *YWHAH* was one of the top genes of this pathway found in both discovery sample and NEWMEDS. *YWHAH* codes for the η subtype of the 14-3-3 protein family, it is expressed mainly in the brain, and it is a positive regulator in the glucocorticoid signal pathway by blocking the degradation of the GR. Variants in this genes have been associated with bipolar disorder and schizophrenia (75).

### 4.2 Limitations

The limitations of the present study should be considered. First, the samples included in this study (particularly NEWMEDS samples used for replication) were heterogeneous from several points of view, for example baseline severity, scales used to assess depressive symptoms, time points of evaluation, antidepressant treatment and dose, setting of recruitment. Heterogeneity across samples and small size of replication samples have limited the power to replicate our findings. In GENDEP and part of STAR*D exome genotypes were available (Illumina Infinium Exome-24 v1.0 BeadChip) but these were not genotyped in the replication samples, making it difficult to replicate the gene and pathway analysis findings. Antidepressant response is known to be a heterogenous phenotype that is affected by a number of genetic and non-genetic variables. In this study we considered age, baseline severity, recruitment centre and ancestry-informative principal components as covariates and we investigated genes and pathways as analysis units in addition to individual SNPs, but there are presumably a number of factors that we did not take into account, such as the effect of the environment. Finally, the replication of findings was weak or absent and no validation of the results was obtained through the use of complementary investigation strategies such as gene expression studies.

### 4.3 Conclusions

The increase in genetic variant coverage seems useful to identify new variants that may influence antidepressant efficacy, but the difficulty in signal replication across different samples is still a problem. Adequate sample size represents a primary issue to allow replication, but also the use of standardized criteria for patient inclusion, treatment and evaluation. Results of pathway analysis and more in general gene set analysis may be easier to replicate across different samples than individual SNPs because sources of heterogeneity or bias (e.g. genotyping or imputation errors) are expected to have a lower influence and effect sizes are expected to be higher than for individual SNPs.

## Acknowledgments

We thank the NIMH for having had the possibility of analyzing their data on the STAR-D sample. We also thank the authors of previous publications in this dataset, and foremost, we thank the patients and their families who accepted to be enrolled in the study. Data and biomaterials were obtained from the limited access datasets distributed from the NIH-supported ‘‘Sequenced Treatment Alternatives to Relieve Depression’’ (STAR*D). The study was supported by NIMH Contract No. N01MH90003 to the University of Texas Southwestern Medical Center. The ClinicalTrials.gov identifier is NCT00021528.

This paper represents independent research funded by the National Institute for Health Research (NIHR) Biomedical Research Centre at South London and Maudsley NHS Foundation Trust and King’s College London. The views expressed are those of the authors and not necessarily those of the NHS, the NIHR, or the Department of Health. The GENDEP project was supported by a European Commission Framework 6 grant (contract reference: LSHB-CT-2003-503428). The Medical Research Council, United Kingdom, and GlaxoSmithKline (G0701420) provided support for genotyping.

The NEWMEDS study was funded by the Innovative Medicine Initiative Joint Undertaking (IMI-JU) under grant agreement n° 115008 of which resources are composed of European Union and the European Federation of Pharmaceutical Industries and Associations (EFPIA) in-kind contribution and financial contribution from the European Union’s Seventh Framework Programme (FP7/2007-2013). EFPIA members Pfizer, Glaxo Smith Kline, and F. Hoffmann La-Roche have contributed work and samples to the project presented here.

The funders had no role in study design, data collection and analysis, decision to publish, or preparation of the manuscript.

The PGRN-AMPS dataset used for the analyses described in this manuscript was obtained from dbGaP (study accession phs000670.v1.p1). PGRN-AMPS was supported, in part, by NIH grants RO1 GM28157, U19 GM61388 (The Pharmacogenomics Research Network), U01 HG005137, R01 CA138461, P20 1P20AA017830-01 (The Mayo Clinic Center for Individualized Treatment of Alcohol Dependence), and a PhRMA Foundation Center of Excellence in Clinical Pharmacology Award.

Dr Rudolf Uher is supported by the Canada Research Chairs Program.

